# WarpDemuX-tRNA: barcode multiplexing for nanopore tRNA sequencing

**DOI:** 10.1101/2025.03.21.644602

**Authors:** Wiep van der Toorn, Isabel S. Naarmann-de Vries, Wang Liu-Wei, Christoph Dieterich, Max von Kleist

## Abstract

Transfer RNA (tRNA) plays an essential role in protein translation and tRNA modifications are important to their function. Recently, nanopore direct RNA sequencing (dRNA-seq) has shown promising results in the detection of complex tRNA modifications. However, its wider adoption in the tRNA field has been limited by a lack of (de)multiplexing solutions. Here, we present WarpDemuX-tRNA: an extension to the WarpDemuX method specifically optimized for multiplexed nanopore tRNA sequencing. Using consensus-based signal analysis using (soft) Dynamic Time Warping and barycenter averaging, our approach improves barcode feature generation and achieves more robust barcode identification. WarpDemuX-tRNA outperforms the original method and achieves 99% precision and 95% recovery for four barcodes, while reducing computational complexity and runtime to six minutes per one million reads. WarpDemuX-tRNA is an open-source and free-to-use solution to high-throughput nanopore tRNA sequencing, facilitating more accessible, cost-effective and high-throughput studies of tRNA modifications and their regulatory mechanisms.

## Introduction

Transfer RNAs (tRNAs) are essential molecules in protein synthesis across all domains of life. Their functionality is intricately regulated through post-transcriptional modifications of specific nucleotides, with more than 100 distinct potential chemical modifications identified to date [Cappannini et al., 2024, Machnicka et al., 2014]. Disruptions in these tRNA modifications are increasingly recognized as causative factors in various human diseases, collectively termed “RNA modopathies” [Suzuki, 2021, Yared et al., 2024, Wang and Lin, 2023, Asano et al., 2018]. These conditions underscore the critical need for scalable, high-resolution methods to profile tRNA modifications, both for understanding disease mechanisms and potential diagnostic applications.

Nanopore direct RNA sequencing (dRNA-seq) has emerged as a powerful technology for studying RNA modifications at single-molecule resolution [Garalde et al., 2018, Workman et al., 2019]. It enables the sequencing of native RNA molecules by measuring ionic current changes as they pass through a biological nanopore. A motor protein is used to guide the RNA through the pore, moving it one nucleotide at a time. At each step, or “event”, a stretch of nucleotides occupying the pore (5-9 bases; “k-mer”) creates a unique electrical signal pattern, with each event generating multiple measurements [Garalde et al., 2018]. The resulting “raw signal” trace provides a direct readout of the molecule’s chemical structure and can be used to simultaneously detect both the sequence of canonical nucleotides and their modifications. This capability makes nanopore technology particularly valuable for studying the epitranscriptome [White and Hesselberth, 2022, Begik et al., 2022, Zhao et al., 2022, Leger et al., 2021, Stoiber et al., 2017].

Although dRNA-seq was initially developed for sequencing long, poly-adenylated RNAs, recent adaptations have enabled the sequencing of short, (non-polyadenylated) tRNAs [Thomas et al., 2021, Lucas et al., 2024]. These adaptations involve the ligation of adapter molecules (“splint adapters”) to both 5’ and 3’ ends of mature tRNAs. The splint adapters effectively lengthen the molecules while also providing a stable ligation site for the standard sequencing adapter required for dRNA-seq. Optionally, after double ligation, the tRNA structure can be removed by linearising the input molecules using reverse transcription, which has been shown to further improve the sequencing yield [Lucas et al., 2024].

Despite these advances in nanopore tRNA sequencing, the lack of open-source (de)multiplexing solutions specific to tRNA sequencing limit the feasibility of large-scale sequencing studies as well as the broader adoption of the technology. Multiplexing reduces the cost of sequencing by allowing the simultaneous sequencing of multiple samples, which is achieved by incorporating unique identifier sequences (barcodes) into each sample prior to sequencing. In nanopore (t)RNA sequencing, the modular design of the standard sequencing adapter, which is a DNA molecule that consists of a constant region containing the pre-loaded motor protein (RMX) and a customizable region (RTA), provides an ideal location for these barcodes. Two community-developed solutions, SeqTagger and WarpDemuX (WDX), have successfully implemented this approach for the latest sequencing chemistry (SQK-RNA004) [Pryszcz et al., 2025, Toorn et al., 2024]. Both methods embed unique DNA-based barcode sequences at the 5’ end of the RTA region, enabling efficient sample multiplexing without modifying the standard library preparation protocol or adding costs.

Notably, these methods were designed for multiplexing of mid-to-long poly-adenylated RNAs and do not account for the unique challenges of nanopore tRNA sequencing. Specifically, they are not optimized for the absence of a poly(A) tail in favor of the 3’ splint adapter that flanks, and differently affects, the barcode signal, nor for the distinct signal characteristics of short and highly modified RNAs like tRNA. These tRNA-specific features require specialized adaptations to ensure reliable barcode detection and accurate demultiplexing. To this day, an open-source and free-to-use multiplexing solution for nanopore tRNA sequencing remains absent from the field. Here, we present WarpDemuX-tRNA (WDX-tRNA), an open-source multiplexing method specifically optimized for nanopore tRNA sequencing with the SQK-RNA004 protocol. Similarly to WDX, WDX-tRNA embeds barcodes at the 5’ end of the RTA adapter and employs Dynamic Time Warping (DTW) for signal analysis. DTW is a dynamic programming algorithm originally developed for speech recognition [Sakoe, 1971, Sakoe and Chiba, 1978], which measures similarity between temporal sequences by finding an optimal alignment through non-linear warping of the time axis, making it robust to variations in speed and timing that are relevant to nanopore signal analysis. In this work, we also employ a variant of DTW called Soft-DTW, which uses a differentiable operator instead of the min operator used in standard DTW, allowing for the use of gradient-based optimization to find the temporal alignment [Cuturi and Blondel, 2018]. Essentially, soft-DTW considers the soft-minimum of the distribution of all costs spanned by all possible alignments between two time series. Soft-DTW is particularly useful when computing time series consensus signals, such as in barycenter averaging [Petitjean et al., 2011, 2014], because such consensus signals can be efficiently learned by gradient descent. Moreover, by considering all possible alignments rather than just the optimal one, soft-DTW has a natural denoising effect which may lead to more robust consensus-based signal analysis [Cuturi, 2011]. In WDX-tRNA, we use soft Dynamic Time Warping Barycenter Averaging (soft-DBA) to generate a consensus signal for the constant sequence of the sequencing adapter (up to the barcode region), which we then use to generate a more precise representation of the barcode signals (“barcode fingerprint”). This improvement enables direct classification of barcode fingerprints, without the need for intermediary fingerprint-to-fingerprint DTW calculations required in the original approach, and thereby reduces the computational cost and time of demultiplexing. The resulting framework enables fast, robust and cost-effective multiplexing of tRNA sequencing experiments at 99% precision and 95% recovery in 6 minutes per million reads, making high-throughput nanopore tRNA sequencing more accessible for the broader research community.

## Results

### Concurrent sequencing of native tRNA populations

To develop and validate our method with known ground truths, we generated two complex datasets of RTA-barcoded mature tRNAs from four species: *H. sapiens, E. coli, C. cerevisiae*, and *C. elegans*, each assigned an unique barcode (**Figure 1a**). Library preparation combined the Nano-tRNA-seq and WDX protocols [Lucas et al., 2024, Toorn et al., 2024], where we replaced the standard sequencing adapter with WDX-barcoded sequencing adapters (**Figure 1b**). The barcoded samples were sequenced using the SQK-RNA004 protocol, with one run per sample. After basecalling, reads were mapped to species-specific tRNA references to establish ground truth labels for evaluating demultiplexing accuracy.

**Fig. 1.**
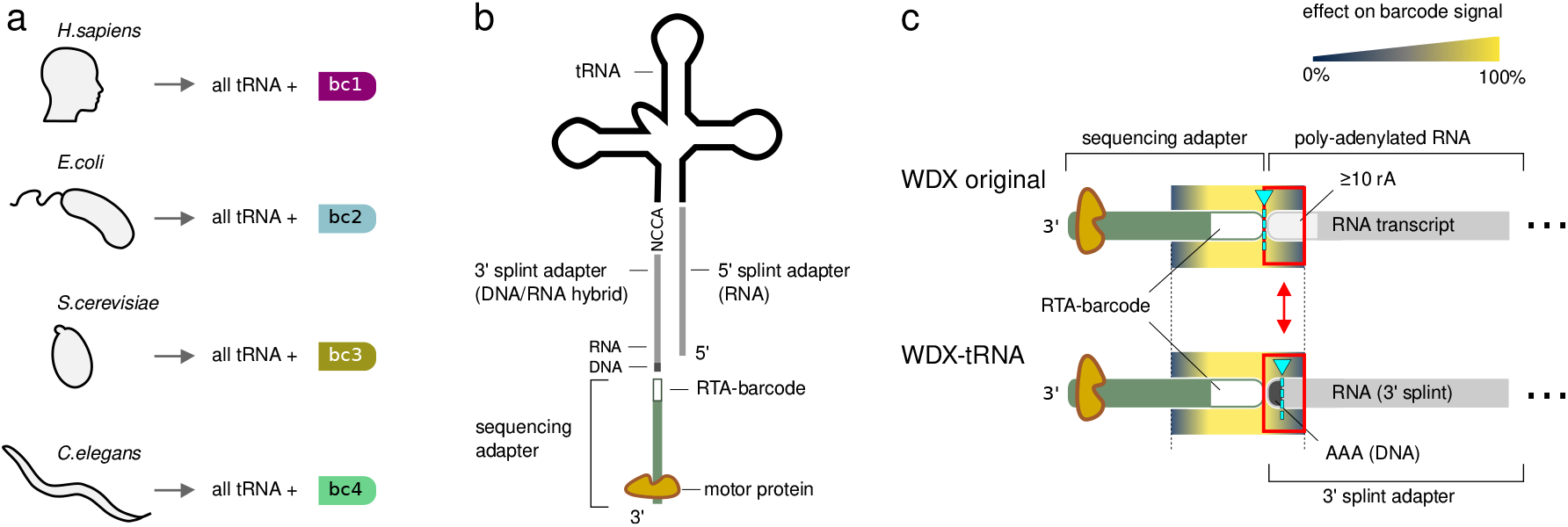
Overview of multiplexed nanopore tRNA sequencing. **a** Experimental design showing the multiplexing of tRNA samples from four species (*H. sapiens, E. coli, C. cerevisiae*, and *C. elegans*), each assigned a unique barcode. **b** Result of library preparation combining Nano-tRNA-seq and WarpDemuX protocols, using barcoded RTA adapters: tRNA (black), 3’ and 5’ splint adapters (grey; with DNA (dark grey) and RNA (light grey) parts of the 3’ splint adapter annotated), standard dRNA-seq sequencing adapter (green) and RTA-barcode region (dark green). **c** Molecular context differences between standard dRNA-seq (polyA tail) and tRNA-specific (splint adapter) protocols, highlighting the altered sequence context of the barcode signal. The DNA-RNA signal boundary which WarpDemuX uses to detect the barcode end is highlighted in blue (triangle, dashed line).

Both datasets achieved similar sequencing depth, with approximately 30-40,000 reads per barcode (**Table S1**). To prevent species-level overfitting and ensure rigorous evaluation, we shuffled the species-to-barcode mappings between the two datasets. Dataset 1 was partitioned into training (42%), validation (14%), calibration (14%), and test (30%, “Test set 1”) sets, while Dataset 2 was reserved entirely for independent testing (“Test set 2”). The heterogeneous nature of these tRNA populations, drawn from diverse species, provided a realistic evaluation of our method’s real-world performance.

A key challenge in adapting WDX for tRNA sequencing stems from differences in the molecular context of the barcode. Due to the nature of nanopore sequencing, the sequence directly surrounding the barcode is known to affect the barcode signal [Liu-Wei et al., 2024]. In both standard dRNA-seq and tRNA sequencing, WDX embeds barcodes at the 5’ end of the RTA adapter.

However, the sequence context of these barcodes differs between protocols: in standard dRNA-seq, the barcode is adjacent to a stretch of RNA adenine bases (also known as the polyA tail), while in tRNA sequencing under the Nano-tRNA-seq protocol, the barcode is flanked by the DNA:RNA hybrid 3’ splint adapter (**Figure 1c**), which, when read in 3’ to 5’ direction as during sequencing, consists of three initial DNA adenine bases followed by an RNA sequence. This altered molecular context of the barcode not only affects the barcode signal characteristics but also changes the DNA-to-RNA transition signal that WDX uses for barcode boundary identification. These tRNA-specific challenges required us to develop new strategies for both signal extraction and analysis.

### Improved Barcode Signal Extraction and Fingerprinting

To address the unique characteristics of tRNA barcode signals, we developed an enhanced algorithm by optimizing the original WDX approach. The original WDX method generates barcode fingerprints through a simple approach: it takes the last *m* segmented events (hyperparameter, default value 25) from the adapter signal. WDX-tRNA improves upon this by separating the fingerprint extraction process into two distinct steps: 1) precise determination of the barcode boundaries by leveraging a consensus signal of the constant adapter sequence preceding the barcode (“adapter header”), followed by 2) a refined segmentation of the identified barcode signal (**Figure 2a**). This two-step approach generates more discriminative barcode fingerprints that are better suited for tRNA sequencing data.

**Fig. 2.**
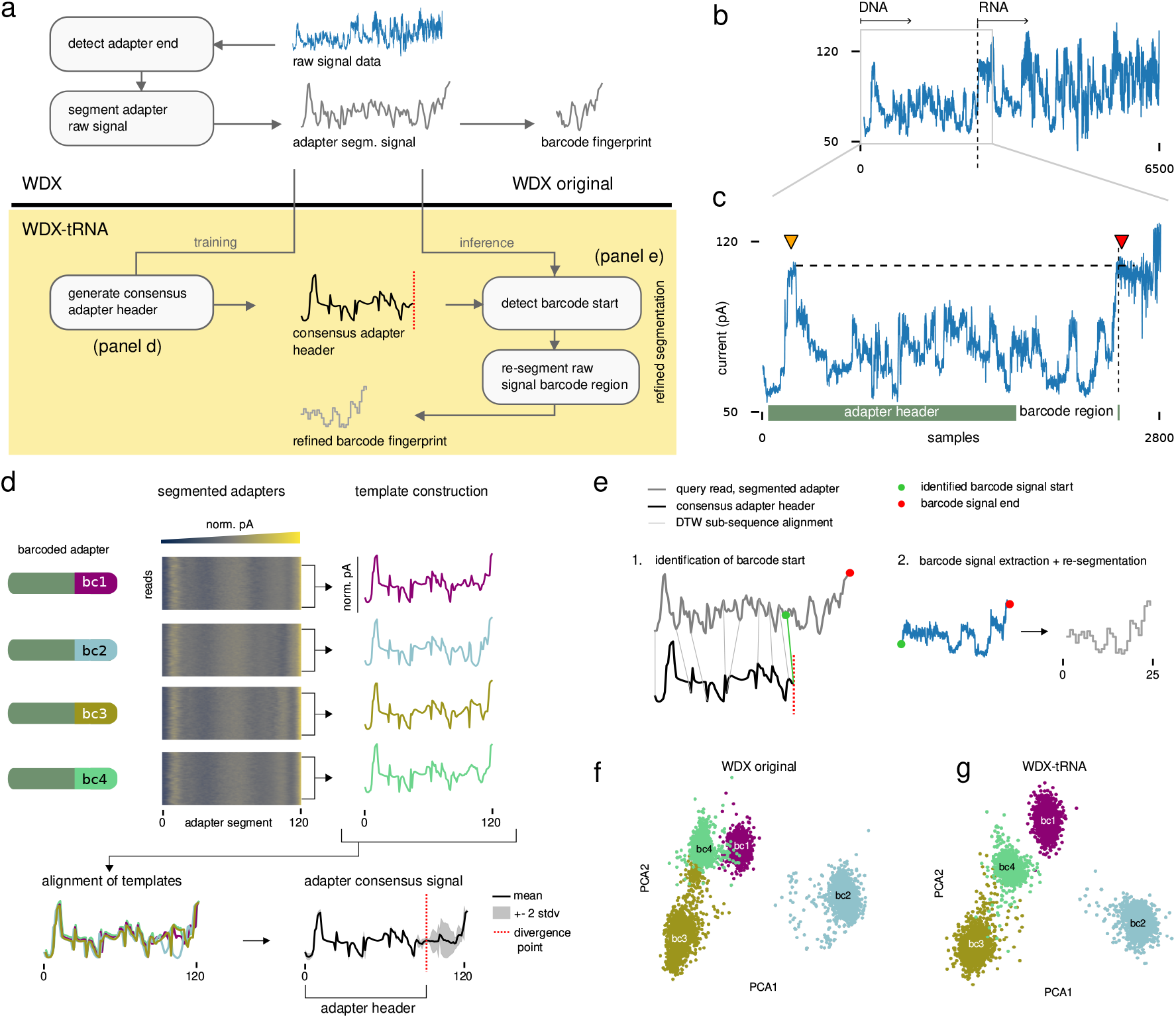
Consensus-based barcode detection workflow. **a** Workflow overview highlighting the common steps (white) and differences (yellow) between the original WarpDemuX and the consensus-based WDX-tRNA methods. **b** Representative nanopore tRNA sequencing raw signal trace. **c** Zoom-in of representative sequencing adapter signal trace. Arrows annotate characteristic high-amplitude signals at the start of the trace (yellow) and at the transition between the DNA and RNA regions (red) used for detection of the barcode end position. **d** Generation of adapter template signals (colored lines) for each barcoded adapter from segmented reads and their alignment to infer the overall adapter consensus (black). The divergence point 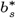 (red dashed line) marks the end of the constant adapter sequence (adapter header) where barcode-specific signals begin to differ from the consensus. **e** Refined barcode segmentation by re-segmentation of barcode signal region after inferring start of region via DTW-subsequence alignment of the adapter header consensus signal. **f-g** Principal Component Analysis (PCA) of barcode fingerprints shows clear separation between different barcode classes (colors), demonstrating the improved discriminative power of WDX-tRNA (g) for tRNA data compared to the original WarpDemuX method (f).

The first critical step in demultiplexing is to precisely locate each barcode’s signal within the raw current trace. This requires accurate identification of both the start and end positions of the barcode signal, a task complicated by the inherent temporal variability of nanopore sequencing. For efficient barcode detection in the original WDX method, barcodes were strategically positioned at the 5’ end of the RTA, where the barcode signal’s end aligned with a distinctive DNA-to-RNA transition point in the signal trace. In the Nano-tRNA-seq protocol, this transition occurs inside the 3’ splint adapter, after the three DNA adenine bases. The start of the RNA sequence within the splint adapter generates a characteristic high-amplitude signal signature that can be used as a reliable marker for the barcode signal’s end (**Figure 2b-c**).

With the barcode end position established, we developed a consensus-based approach to identify the barcode start position. In both WDX methods, to reduce computational complexity and measurement noise, all adapter signals are segmented before further processing by partitioning them into regions of uniform current levels, using the mean value per region to efficiently down-sample the signal. In WDX-tRNA, we first use these segmented adapter signals to generate template signals for each barcoded adapter using soft-DBA. By aligning the template signals, we obtain an overall adapter consensus. We then identified the divergence point that marks the position where the aligned barcode-specific signals begin to differ (**Figure 2d**). The consensus sequence up to this divergence point represents the constant adapter sequence that precedes each barcode (“adapter header”) that we use to determine the barcode start position in new reads.

For each new read, we determine the barcode start position by a subsequence alignment of the adapter header consensus signal to the segmented adapter signal using DTW, where the end of the alignment is the barcode start position. Using the segmentation event lengths, we can infer the corresponding location in the raw signal trace. With both barcode region boundaries established, we extract the barcode raw signal and perform a refined resegmentation to obtain a more precise barcode fingerprint (see Methods, **Figure 2e**).

The objective of the refined segmentation is to generate barcode fingerprints with greater discriminative power for classification. To evaluate the improvement in fingerprint quality, we assessed their class separability after resolving temporal variations. Temporal alignment was achieved by aligning fingerprints to their respective barcode consensus, calculated using soft-DBA. Principal Component Analysis (PCA) of these temporally resolved fingerprints shows improved separation between different barcode classes, demonstrating improved discriminative potential compared to the original method (**Figure 2f-g**).

### Direct and Efficient Barcode Classification

We developed an efficient classification approach that directly leverages the refined barcode fingerprints. Rather than computing DTW distances between query fingerprints and training samples as in the original WDX method, we train gradient boosted trees directly using the barcode fingerprints as input features. This approach learns to distinguish between different barcode classes from the improved fingerprint patterns, while reducing computational overhead by eliminating the need for DTW distance calculations during inference.

To ensure reliable classification, we assign each prediction a confidence score [Toorn et al., 2024]. Reads with confidence scores below barcode-specific thresholds are filtered out as “unclassified”. These thresholds are calibrated for each barcode independently using a held-out calibration dataset to achieve a target precision level of 99%. Depending on their use-case, users can choose to increase or decrease the precision level by adjusting the thresholds (see **Fig. S1**).

The classifier achieves high performance across all barcodes, with precision and recall (**Figure 3a**) and Receiver Operating Characteristic Area Under the Curve (ROC AUC) scores (**Figure 3b**) around 0.99 while achieving recovery (% of reads assigned) of around 95% on an independent sequencing run (Test set 2, **Table 1**). When comparing the confusion matrices of the original WDX method and WDX-tRNA (**Figure 3c-d**), we found that WDX-tRNA in particular reduced the misclassification rate of barcode 4 to 1, while at the same time substantially increasing the recovery rate of barcode 2-4 (**Figure 3e**).

**Fig. 3.**
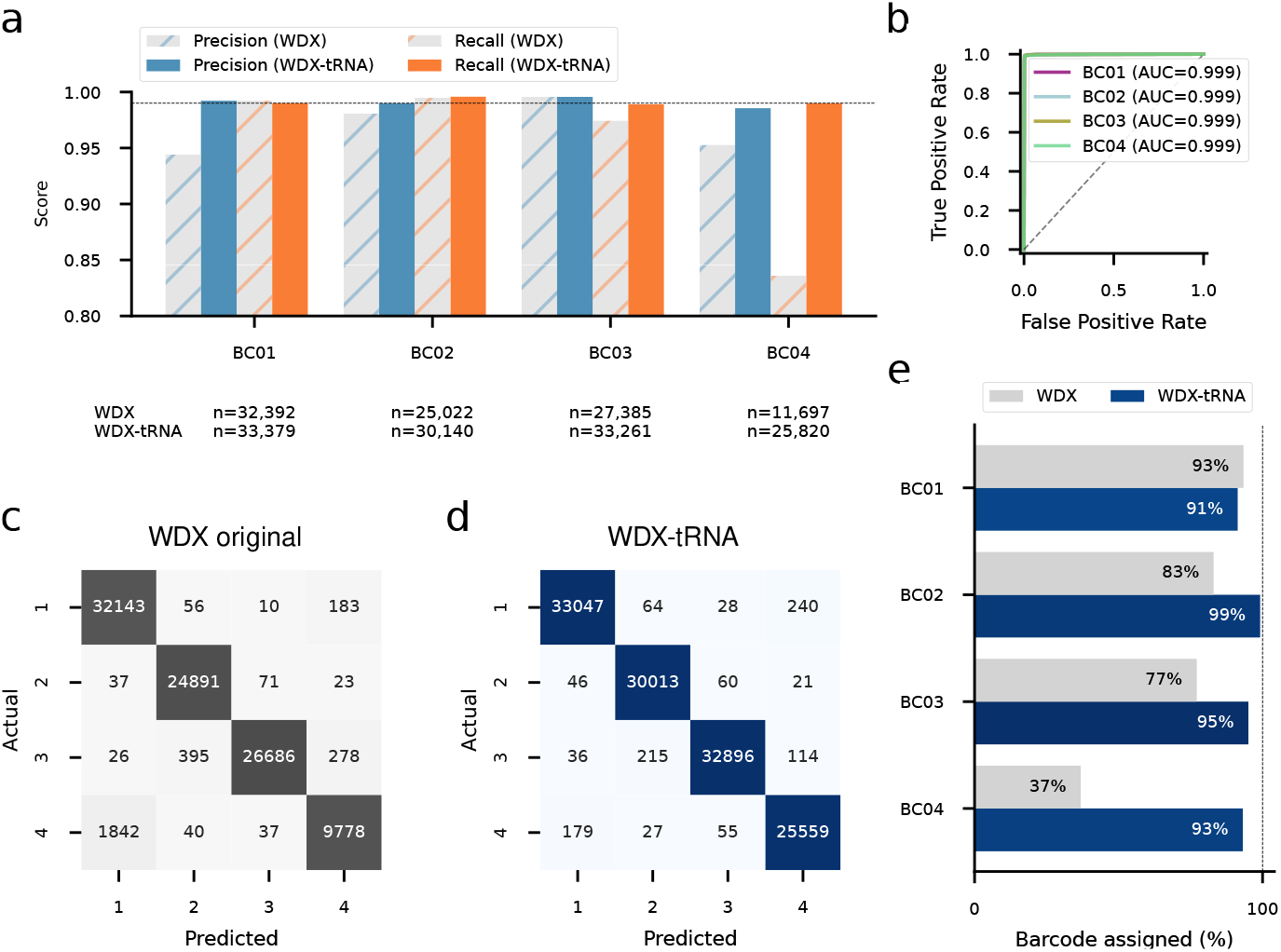
Performance comparison of WDX-tRNA and WDX original classification on independent replicate (Test set 2). **a** Precision (blue) and Recall (orange) metrics for individual barcodes using WDX original (gray hatched) versus WDX-tRNA (solid color). Reference line at 0.99 (black dashed line). **b** Receiver Operating Characteristic (ROC) curve per barcode (colors), substantially outperforming random classification (diagonal gray dashed line). **c-d** Confusion matrices for WDX original (c, gray) and WDX-tRNA (d, blue). **e** Classification performance showing improved yield (% of barcodes assigned) across all barcodes using WDX-tRNA (blue) versus WDX original (gray).

Moreover, even with the added complexity of the refined barcode fingerprinting, the elimination of DTW distance calculations during inference resulted in a 2.76-fold increase in processing speed, enabling the demultiplexing of 100,000 reads in 38 seconds (16 cores, **Table 2**).

**Table 1.** Comparison of classification performance between WarpDemuX-tRNA and original WarpDemuX models on two independent test sets. Values show WarpDemuX-tRNA performance metrics with relative changes compared to original WarpDemuX in parentheses. Relative change is omitted for values less than 1%. Precision and recall values reflect classification on assigned reads, while recovery indicates the percentage of assigned reads (reads passing confidence score filtering). ‘Macro avg’ represents the unweighted mean across all barcodes.

| Dataset | BC | Precision | Recall | Recovery |
| --- | --- | --- | --- | --- |
| Test set 1 | BC 1 | 0.991 (+7%) | 0.992 | 92.83 (+14%) |
|  | BC 2 | 0.990 (+2%) | 0.995 | 98.44 (+32%) |
|  | BC 3 | 0.977 | 0.991 (+2%) | 96.46 (+21%) |
|  | BC 4 | 0.990 (+4%) | 0.965 (+25%) | 92.30 (+131%) |
|  | macro avg | 0.987 (+3%) | 0.986 (+6%) | 95.09 (+37%) |
| Test set 2 | BC 1 | 0.992 (+6%) | 0.990 | 91.54 (+15%) |
|  | BC 2 | 0.990 (+1%) | 0.996 | 98.56 (+33%) |
|  | BC 3 | 0.996 | 0.989 (+1%) | 95.72 (+22%) |
|  | BC 4 | 0.986 (+4%) | 0.990 (+20%) | 92.74 (+126%) |
|  | macro avg | 0.991 (+3%) | 0.991 (+5%) | 94.57 (+36%) |

**Table 2.** Run time comparison of WarpDemuX-tRNA fingerprint and WarpDemuX DTW-SVM demultiplexing. Runtime benchmarks were executed on a high-performance server using different numbers of Intel(R) Xeon(R) Gold 6130 CPUs @ 2.10GHz (#CPU) and 24GB RAM. Time displayed as minutes (MM) and seconds (SS), with mean and standard deviation (std) over 5 runs.

| Model | #CPU | Time (MM:SS) |  |  |  | Relative difference |
| --- | --- | --- | --- | --- | --- | --- |
|  |  | 100,000 reads |  | 1 million reads |  |  |
|  |  | mean | std | mean | std |  |
| WDX original | 8 | 02:31 | 00:00 | 25:12 | 00:00 | — |
|  | 16 | 01:45 | 00:04 | 17:32 | 00:47 | — |
|  | 24 | 01:19 | 00:00 | 13:12 | 00:08 | — |
| WDX-tRNA | 8 | 01:04 | 00:00 | 10:49 | 00:03 | -57.1% |
|  | 16 | 00:38 | 00:00 | 06:20 | 00:03 | -63.9% |
|  | 24 | 00:36 | 00:01 | 06:07 | 00:13 | -53.6% |

## Discussion

The ability to multiplex nanopore tRNA sequencing samples is crucial for making nanopore tRNA sequencing economically viable for large-scale studies, such as modification profiling. Our enhanced demultiplexing approach addresses key challenges specific to tRNA sequencing while maintaining the simplicity and cost-effectiveness of barcode incorporation through the RTA adapter.

The improved performance of our method stems from three key innovations. First, our consensus-based approach to signal boundary detection proves more robust to the inherent variability in tRNA sequencing signals. Unlike mid-to-long RNA sequencing, where the longer molecule length helps stabilize translocation speed and facilitates signal normalization, tRNA sequencing produces very short reads that exhibit greater signal variability. Our method effectively handles these challenges by leveraging collective signal patterns across multiple reads to establish reliable barcode boundaries. Second, our two-step extraction process, in which the identified barcode signal region is subjected to a focused re-segmentation, generates more discriminative barcode fingerprints. By first identifying the exact barcode region using the adapter header consensus and then performing a focused re-segmentation, we capture the distinctive characteristics of each barcode more accurately. Third, the improved quality of these fingerprints enables direct classification using gradient boosting trees, eliminating the computationally expensive DTW distance calculations during inference that were required in the original method. This not only speeds up demultiplexing but also improves classification accuracy by allowing the model to learn complex patterns directly from the refined fingerprint features.

Both WDX and WDX-tRNA rely on signal segmentation as a fundamental processing step and employ a computationally efficient segmentation algorithm that achieves fast processing speeds by making locally optimal choices at each step (“greedy”). While this approach enables rapid processing, it may result in sub-optimal segmentation results, particularly for longer signals where segmentation errors can compound due to the sequential nature of greedy algorithms. To mitigate potential information loss due to segmentation errors, both methods deliberately over-segments adapter signals relative to the known number of events based on the sequence. This conservative strategy reduces the risk of merging distinct sequencing events into a single segment, which would result in loss of signal characteristics. However, it also means there is no guaranteed one-to-one correspondence between segmented signals from different reads, necessitating the use of temporally robust and computationally expensive distance metrics (such as the DTW distance) when comparing them.

WDX-tRNA addresses this limitation by introducing a novel two-step segmentation strategy to obtain barcode fingerprints that are directly usable for classification without the need for temporally robust measures. We first establish precise barcode boundaries from oversegmented adapter signals through consensus-alignment before performing a focused re-segmentation of the shorter barcode region. The alignment step serves a dual purpose: it not only identifies the barcode position but also functions as a quality control step. Reads that fail to produce alignments within calibrated bounds are flagged as potentially problematic (e.g., due to adapter end misidentification, blocked pores, concatemers, or segmentation errors) and excluded from further analysis. After successful barcode position identification, the second segmentation phase benefits from analyzing a substantially shorter signal with fewer breakpoints (barcode-only versus full adapter), which reduces the effect of compounding inaccuracies inherent to greedy segmentation. The combination of precise boundary detection and focused re-segmentation produces more discriminative barcode fingerprints, as evidenced by the improved class separation (**Figure 2f-g**).

While our method achieves high classification performance overall, our analysis reveals important insights about barcode signal characteristics in the tRNA sequencing context. Notably, the ionic current signals generated by different barcode sequences exhibit varying degrees of distinctiveness. In particular, barcodes 1 and 4, and barcodes 3 and 4 produce more similar signal patterns, as evidenced by their proximity in the dimensionality-reduced space (**Figure 2f-g**). This reduced separability, compared to the level observed in the original WDX work where they are used within the dRNA-seq context, may be attributed to a structural difference in the tRNA protocol: the presence of the three additional DNA adenine bases at the start of the splint adapter. WDX barcodes were optimized for the standard RTA-polyA setup used in mRNA sequencing, where the barcode signal is directly flanked by RNA adenine basses (polyA tail). The additional DNA bases in the tRNA protocol affect the end of the barcode signal and may reduce their distinctiveness under the present analysis.

Despite these challenges, our method successfully distinguishes between barcode signals. However, future improvements could focus on optimizing barcode sequences to maximize signal distinctiveness in the context of the splint adapter. By selecting sequences that generate more divergent ionic current patterns under the tRNA sequencing protocol, we could potentially further enhance classification performance and/or expand the number of available barcodes.

In summary, our enhanced demultiplexing approach represents a significant advancement for multiplexed nanopore tRNA sequencing. By combining robust signal processing with efficient classification, we enable reliable sample identification for up to four samples while reducing computational overhead. The availability of this open-source solution removes commercial barriers for multiplexed nanopore tRNA sequencing, while enabling community-driven improvements and adaptations, fostering broader adoption of nanopore sequencing in tRNA research.

## Methods

### Sample preparation and tRNA extraction

*H. sapiens* - Total HEK293 (DSMZ, ACC 305) RNA was isolated with Trizol (Thermo Fisher Scientific) according to the manufacturer’s instructions. *C. elegans* - Total RNA was a kind gift from Oliver Hoppe and Carmen Nussbaum (Bukau Lab). *S. cerevisiae* - Total RNA was a kind gift from Katja Strässer. *E. coli* - Total RNA was a kind gift from Frederik Weber and Andres Jäschke. Small and large RNAs were separated using the RNA Clean and Concentrator-25 kit (Zymo Research) with the respective protocol. 4 µg small RNA were deacylated by incubation in 100 mM Tris pH 9.0 for 30 min at 37°C and purified afterwards with RNA Clean and Concentrator-5 kit (Zymo Research).

### Library preparation for Nanopore Sequencing

Equal amounts (200 to 500 ng) of deacylated tRNAs were subjected to splint adapter ligation with T4 RNA ligase 2 as described [Lucas et al., 2024]. Reactions were cleaned up with 1.8x BioDynami tRNA SPRI beads (Nucleus Biotech) and eluted in 10 µl H_2_O. The complete splinted tRNAs were ligated to WDX-barcoded RTA adapters (WDX-RTA 4, 5, 7, and 11, see **Table S2**) with T4 DNA ligase as described [Lucas et al., 2024]. The species to barcode mapping per dataset is shown in **Table S1**. Ligation products were reverse transcribed employing Induoro reverse transcriptase (30 min, 60°C, followed by 10 min 70°C). Reactions were stopped by addition of EDTA to a final concentration of 250 mM and cleaned up with 1.8x BioDynami tRNA SPRI beads and eluted in 10 µl H_2_O. 5 µl of each sample were ligated to the RLA adapter as described [Lucas et al., 2024] followed by a final clean up step with BioDynami SPRI beads.

### Nanopore sequencing

Samples were sequenced on a P24 A-series device using FLO-PRO004RA flow cells (Oxford Nanopore Technologies). After priming the flow cell, the prepared sequencing libraries were loaded completely onto the flow cells. Sequencing was performed using live basecalling with Dorado Basecall Server 7.4.13 (MinKNOW 24.06.14), using the High Accuracy model (HAC), with a Q score threshold of 7.

### Data Processing

Our set of tRNA samples from 4 species covers a long phylogenetic distance. Each species is assigned to a specific barcode, which we can validate by alignment to a reference database of nuclear-encoded RNA sequences for all 4 species. We compiled a database of reference sequences from gtrnadb.ucsc.edu. All tRNA reads were aligned to this reference using Parasail (version 2.6.0), an optimal local sequence aligner, combined with a robust statistical filtering approach. To ensure high-confidence alignments, we use a simulation-based significance assessment that distinguishes true alignments from random matches by comparing forward alignments against a null distribution generated from reverse sequence alignments, as proposed by Sun et al. [2023]. We filtered the resulting alignments using a 99.9% specificity threshold.

### Adapter terminus detection

In nanopore tRNA sequencing using the Nano-tRNA-seq protocol, the barcode signal is flanked by a distinctive transition where the DNA sequencing adapter meets the 3’ DNA:RNA hybrid splint adapter. The structure of the splint adapter’s 3’ tail—three DNA adenine bases followed by 10 rA and two rG bases (read in 3’ to 5’ direction as during sequencing)—creates a characteristic high-amplitude signal pattern that serves as a reliable marker for the barcode signal’s end.

To efficiently detect this transition, we extract the first 10,000 samples of the raw signal and down-sample it by a factor of 10. The adapter signal contains two key features: a characteristic high-amplitude peak at the start of the read, followed by a second peak marking the transition to the splint adapter. Both peaks are highly conserved across reads.

We identify the maximum current peak in the first 150 samples of the down-sampled signal, then search for the next peak that exceeds this maximum, starting from a fixed offset. The timing between these peaks and their relative amplitudes are used to identify the adapter terminus and flag potential problematic reads.

### Initial segmentation

The segmentation process computes a series of *n* − 1 breakpoints that define *n* regions of relatively uniform signal levels. We use *n* = 120 and take the mean per region to get a signal of length *n* per barcode, efficiency down-sampling the signal and reducing measurement noise, both of which reduce the complexity of the following steps.

To detect signal transitions, we use a sliding window approach that compares the statistical properties between adjacent windows. For each position *t* in the signal, we compute a discrepancy measure *d* (here: the absolute value of the t-statistic) between two windows of length *w*, positioned before and after *t*. The discrepancy is low when both windows contain similar signals (within the same segment) and high when they span a transition point. We identify change points by detecting peaks in the resulting discrepancy curve that are at least *l* samples apart. This computationally efficient approach scales linearly with signal length *n* and window size *w* (*O*(*nw*)). We selected *w* = 18 and *l* = 9 following hyper-parameter tuning on the training data.

To account for variability in current levels across reads and signal drift, we normalize signals based on their initial segmentation mean and standard deviation prior to calculating the refined barcode fingerprint (see **Re-segmentation of the barcode signal**).

### Constructing the adapter header consensus signal

After adapter terminus detection, segmentation and normalization (zero-mean, unit-variance per signal), we construct template signals for each barcoded adapter using soft-DBA (as implemented in the tslearn package [Tavenard et al., 2020], *λ* = 0.1). We resample the training data to compute 500 templates per barcoded adapter, each calculated from 100 randomly sampled signals. These template signals are then averaged again using soft-DBA to create an adapter template per barcoded adapter. By aligning the individual adapter templates using DTW, we can identify the barcode start position as the location where the template signals begin to diverge. The adapter header consensus signal is the mean of the aligned template signals up to the divergence point.

### Re-segmentation of the barcode signal

For a given read, we first determine its barcode start position by DTW subsequence alignment of the consensus adapter header to its segmented adapter signal. Subsequence alignment results in a start and end position of the consensus adapter header in the read’s adapter segment signal. We establish lower and upper bounds for each position as the 1st and 99th percentiles of the training data distribution, respectively, but set the lower bound of the start position to −1 (no filtering). Reads with alignment positions outside these bounds are discarded (**Fig. S2**). For the remaining reads, we map the end of the alignment back to the read’s raw signal to identify the start of the barcode signal region. Using the previously determined adapter terminus position (end of barcode region), we extract the barcode signal and perform a refined segmentation using the windowed approach previously described, with *n* = 25 breakpoints, window size *w* = 18 and minimal event length *l* = 9, and use the last *m* = 25 segments. The resulting barcode fingerprint is standard-normalized with respect to the mean and standard deviation of the initial adapter segmentation.

### Barcode classification

We use gradient boosting trees (as implemented in CatBoost [Prokhorenkova et al., 2019]) for barcode classification, which builds an ensemble of weak decision trees in an iterative fashion to create a strong classifier. Each tree attempts to correct the errors made by the previous trees in the ensemble. For multiclass classification with *K* classes, CatBoost uses the multiclass cross-entropy loss function:

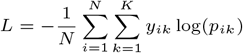

where *N* is the number of samples, *y*_*ik*_ is a binary indicator whether class label *k* is the correct classification for sample *i*, and *p*_*ik*_ is the predicted probability that sample *i* belongs to class *k*.

The model is trained on the refined barcode fingerprints using 2000 iterations with early stopping after 20 rounds without improvement on the validation set. We use a learning rate of 0.1 and a maximum tree depth of 6 to control model complexity. To further prevent overfitting, we set a minimum of 5 samples required in each leaf node. The predicted probabilities for each class are obtained through a softmax transformation of the raw model outputs:

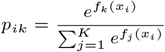

where *f*_*k*_(*x*_*i*_) is the raw model output for class *k* on sample *i*.

For each prediction, we compute a confidence score as the difference between the highest and second-highest class probabilities [Toorn et al., 2024]. This margin-based confidence score provides a measure of prediction reliability, with larger margins indicating more confident predictions. We calibrate barcode-specific confidence thresholds on a held-out calibration set to achieve 99% precision, filtering out predictions below these thresholds as “unclassified”.

## Data availability

All raw sequencing data for this study have been deposited in the European Nucleotide Archive (ENA) at EMBL-EBI under accession number PRJEB87328.

## Code availability

Code is available at https://github.com/KleistLab/WarpDemuX (v0.4.7) and via Zonodo [Van der Toorn et al., 2025].

## Acknowledgments

W.L.-W. was funded by the European Union’s Horizon 2020 research and innovation program under the MarieSklowska-Curie Actions Innovative Training Networks grant, agreement no. 955974. M.v.K and W.L.-W. acknowledge funding provided by the German Ministry for Science and Education, grant number 01KI2016. M.v.K. acknowledges funding by the Deutsche Forschungsgemeinschaft (DFG, German Research Foundation) under Germany’s Excellence Strategy—The Berlin Mathematics Research Center MATH+ (EXC-2046/1, project ID: 390685689). C.D. acknowledges funding by the German RMaP consortium (Deutsche Forschungsgemeinschaft [DFG, German Research Foundation, 439669440 TRR319 RMaP TP B01])

## Author contributions

I. N-d.V. performed the wet lab experiments. W. vdT. and C. D. performed the bioinformatic analysis. W. vdT. conceived the work. C. D. and M. vK. supervised the work. C.D. and M. vK. acquired funding to conduct the work. W. vdT. prepared the figures. W.L.-W. provided substantial feedback throughout the development of the work, including figure preparation and paper writing. W. vdT. and I. N-d.V. wrote the paper, with contributions from all authors.

