## Supplementary Information for "WarpDemuX-tRNA: barcode multiplexing for nanopore tRNA sequencing"

### Supplementary Tables

**Table S1.** Dataset characteristics. For each replicate, the number of reads after alignment (aligned) and after barcode fingerprint calculation (fpts) are shown.

| **replicate** | **BC** | **species** | **aligned** | **fpts** | **fpts rate (%)** |
|---|---|---|---|---|---|
| 1 | 1 | *H.sapiens* | 43627 | 40708 | 93.3 |
| | 2 | *E.coli* | 43158 | 39377 | 91.2 |
| | 3 | *S.cerevisiae* | 44466 | 41824 | 94.1 |
| | 4 | *C.elegans* | 38050 | 35463 | 93.2 |
| 2 | 1 | *H.sapiens* | 39256 | 36463 | 92.9 |
| | 2 | *C.elegans* | 32244 | 30580 | 94.8 |
| | 3 | *S.cerevisiae* | 37229 | 34750 | 93.3 |
| | 4 | *E.coli* | 30116 | 27841 | 92.4 |

**Table S2.** Barcode sequences. WDX-RTA barcodes used in this study, sequences in 5' to 3' direction.

| BC | WDX-RTA | Sequence |
|---|---|---|
| 1 | Barcode 4 | GGAGGCCAGGCGGACCGA |
| 2 | Barcode 5 | ACGGACCTTTTGACTTAA |
| 3 | Barcode 7 | CCACGGAGGGAGGATTGG |
| 4 | Barcode 11 | GCCCGCCGGGGGAGAAGC |

### Supplementary Figures

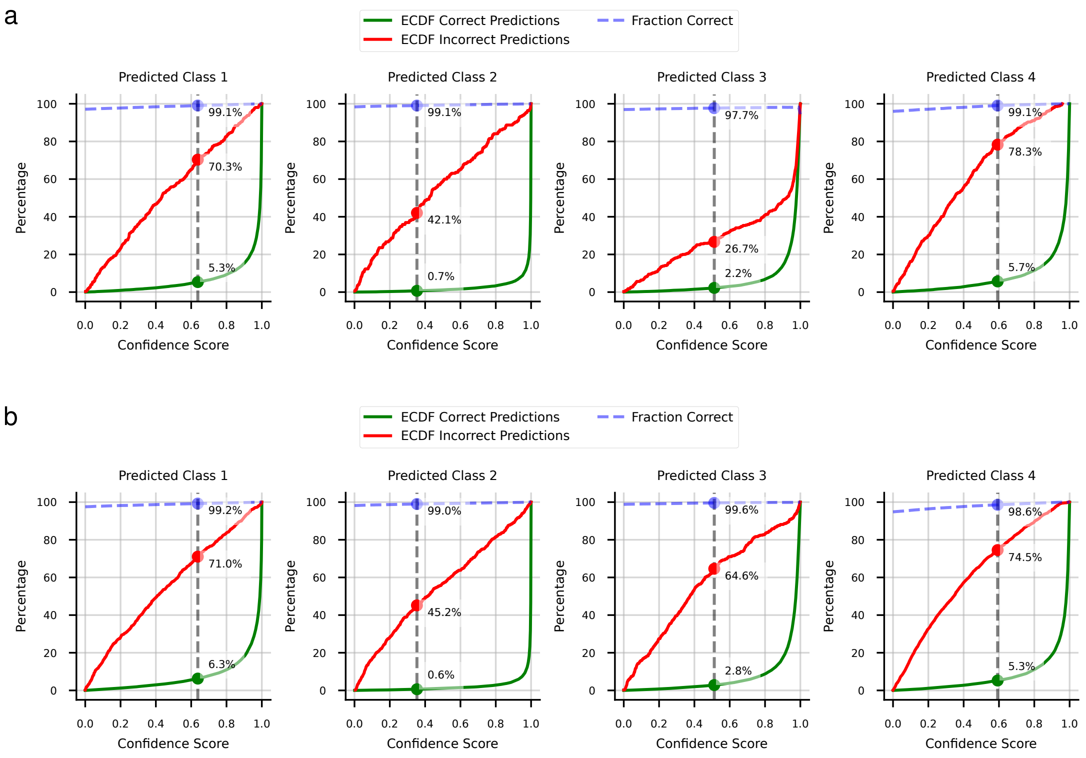


**Fig. S1.** Empirical cumulative distribution functions (ECDFs) showing the relationship between confidence scores and prediction outcomes across four classes. For each predicted class, red lines represent true positive predictions (correct classifications) while green lines represent false positive predictions (incorrect classifications). The vertical dashed gray line indicates the calibrated 99% precision confidence threshold, with corresponding percentages of predictions below this threshold shown for both true positives (red) and false positives (green). The blue dashed line and dot represent the percentage of correct predictions as a function of the confidence score. **a** Test set 1. **b** Test set 2.

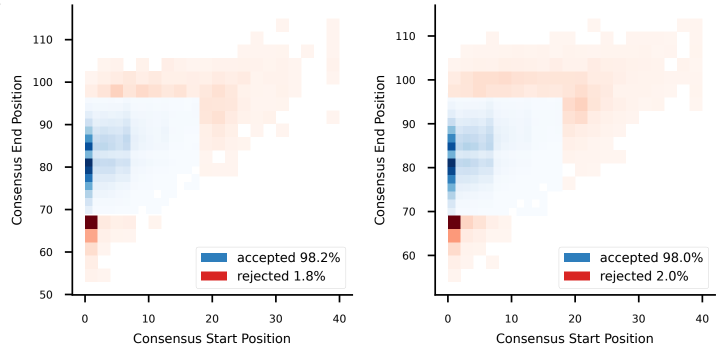


**Fig. S2.** Distribution of consensus subsequence alignment positions. Two-dimensional histogram showing the relationship between the start (x-axis) and end (y-axis) positions of the consensus subsequence alignment per read. Blue shading represents the density of accepted reads while red shading indicates rejected reads (positions not within bounds). **a** Test set 1 (1.82% reads rejected). **b** Test set 2 (2.02% reads rejected).
